# Cyclin B3 is dispensable for mouse spermatogenesis

**DOI:** 10.1101/608315

**Authors:** Mehmet E. Karasu, Scott Keeney

## Abstract

Cyclins, as regulatory partners of cyclin-dependent kinases (CDKs), control the switch-like cell cycle transitions that orchestrate orderly duplication and segregation of genomes. Compared to mitosis, relatively little is known about how cyclin-CDK complexes control meiosis, the specialized cell division that generates gametes for sexual production. Mouse cyclin B3 was previously shown to have expression restricted to the beginning of meiosis, making it a candidate to regulate meiotic events. Indeed, female mice lacking cyclin B3 are sterile because oocytes arrest at the metaphase-to-anaphase transition of meiosis I. However, whether cyclin B3 functions during spermatogenesis was untested. Here, we found that males lacking cyclin B3 are fertile and show no detectable defects in spermatogenesis based on histological analysis of seminiferous tubules. Cytological analysis further showed no detectable defects in homologous chromosome synapsis or meiotic progression, and suggested that recombination is initiated and completed efficiently. Moreover, absence of cyclin B3 did not exacerbate previously described meiotic defects in mutants deficient for cyclin E2, suggesting a lack of redundancy between these cyclins. Thus, unlike in females, cyclin B3 is not essential for meiosis in males despite its prominent meiosis-specific expression.

## Introduction

During meiosis, one round of chromosome replication is followed by two rounds of nuclear division, which allows organisms to halve their genome content. Unique to meiosis, homologous chromosomes segregate in the first division, then sister chromatids are separated in the second division. To achieve faithful homolog separation, programmed DNA double-strand breaks (DSBs), DSB repair via homologous recombination, and formation of chiasmata between homologs must occur (Lam and Keeney 2015).

Homologous recombination is tightly regulated during meiotic prophase I. One layer of regulation is implemented by the cell cycle components, cyclins and cyclin-dependent kinases (CDKs) (Evans et al. 1983). Cyclins share a conserved domain called a “cyclin box”, and are expressed and degraded in ordered fashion during the cell cycle. A well-known function for cyclins is to regulate the activity of their catalytic binding partners, CDKs.

Cyclins with or without their CDK partners regulate various events during meiotic prophase I. For example, in *Saccharomyces cerevisiae*, phosphorylation of Mer2 by cyclin-CDK complexes is essential for the initiation of meiotic recombination (Henderson et al. 2006). In *Tetrahymena thermophila*, the meiosis-specific cyclin Cyc2p is essential for formation of the micronuclear bouquet and for homologous pairing and synapsis (Xu et al. 2019). In *Caenorhabditis elegans*, cyclin-related protein COSA-1 is a key component for the designation of crossover recombination products (Yokoo et al. 2012).

In mammalian meiosis, cyclins are sequentially expressed (reviewed in (Wolgemuth and Roberts 2010; Wolgemuth et al. 2013)) and genetic ablation of cyclin or CDK genes has demonstrated important roles in gametogenesis. For example, disruption of *Cyclin A1* (*Ccna1*), which is expressed in late-prophase spermatocytes but not in somatic cells, causes arrest in late meiotic prophase and infertility in male mice (Liu et al. 1998). Mutation of *Cyclin E1* (*Ccne1*) and *Cyclin E2* (*Ccne2*) causes early meiotic prophase defects (Martinerie et al. 2014). In addition, mice lacking *Cdk2* exhibit a defect in the repair of DSBs and pairing of homologous chromosomes, causing meiotic arrest and infertility (Berthet et al. 2003; Ortega et al. 2003).

Mouse *Cyclin B3* (*Ccnb3*), an X-linked gene, is germ-line specific and is expressed in both spermatocytes and oocytes during the leptotene and zygotene stages of meiotic prophase I (Nguyen et al. 2002). The importance of the restricted expression of *Ccnb3* was tested in a mouse model in which prolonging expression of *Ccnb3* beyond early meiotic prophase caused abnormal spermatogenesis and increased apoptosis (Refik-Rogers et al. 2006). However, whether the absence of *Ccnb3* affects spermatogenesis remained unclear.

Given the confined expression of *Ccnb3*, we hypothesized that *Ccnb3* might regulate events during early meiotic prophase. In this study, we tested this hypothesis by generating and characterizing a mutant *Ccnb3* allele via CRISPR–Cas9-mediated gene targeting. In separate work, we and others found that cyclin B3-deficient females are sterile (Karasu et al. 2019; Li et al. 2019). We show here, in contrast, that male mice lacking *Ccnb3* are fertile and display little or no evidence of meiotic abnormalities. Thus, contrary to expectation, *Ccnb3* is dispensable for spermatogenesis.

## Results

### Cyclin B3 antibody generation and cyclin B3 expression during the first wave of spermatogenesis

The *Ccnb3* locus gives rise to a 4.1 kb gene product, which codes for a 157.9 kDa protein. This is an unusual size for mammalian cyclins, which are typically around 50 kDa (Evans et al. 1983; Bloom and Cross 2007). Until now, although *Ccnb3* mRNA could be detected in mammalian germ cells (Lozano et al. 2002; Nguyen et al. 2002), endogenous cyclin B3 from mouse testis extracts has not been detected due to the lack of a suitable antibody. To overcome this issue, we generated monoclonal antibodies against cyclin B3. A total of 23 monoclonal antibodies were raised against 8 different peptides (**Fig 1a**).

**Figure 1:**
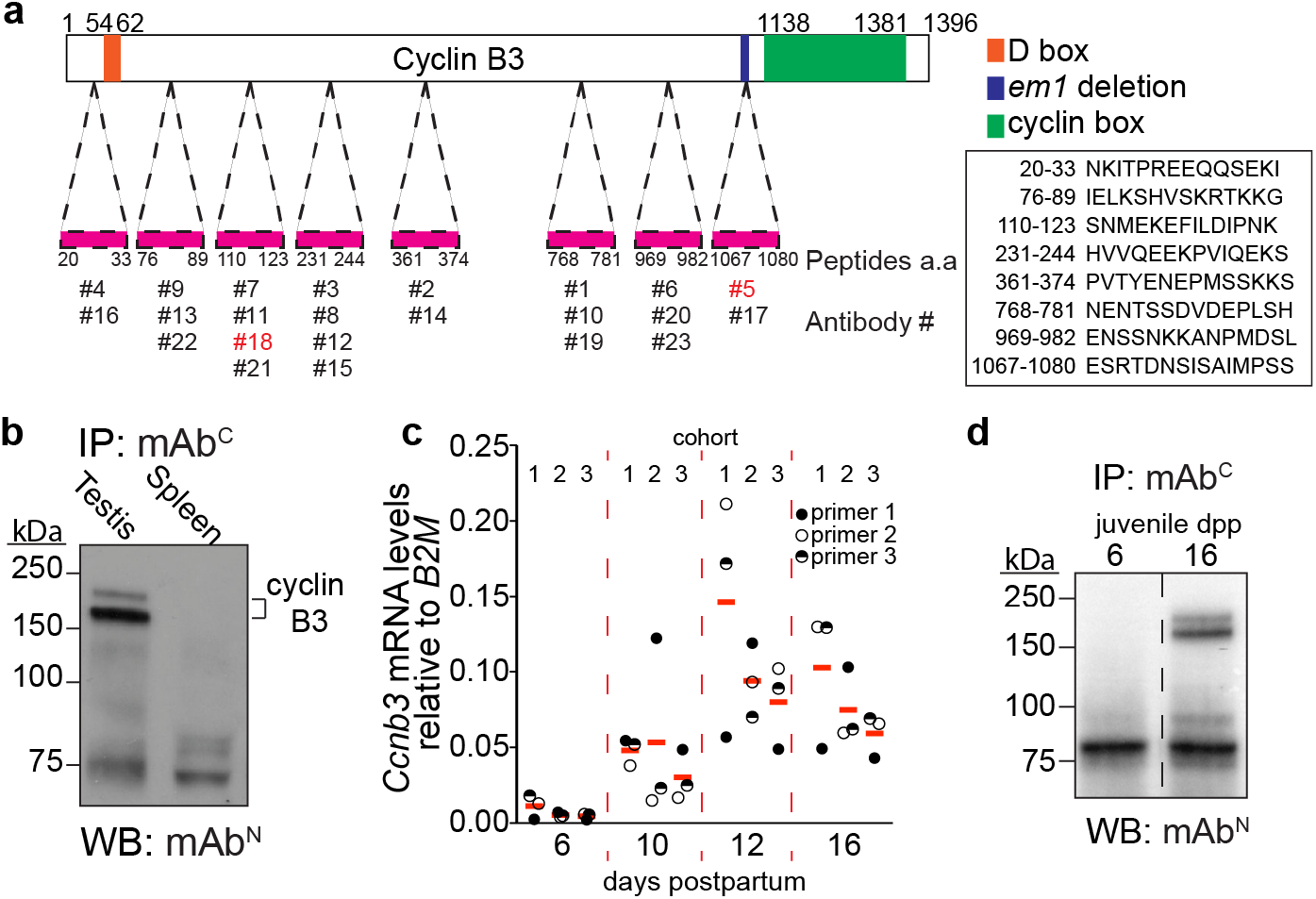
Antibody generation and cyclin B3 expression in the first wave of spermatogenesis. a) Schematic of cyclin B3 showing the cyclin box domain (green) and the destruction box (orange). The positions and sequences of the peptides used to raise antibodies are shown. b) IP-WB for endogenous cyclin B3 in extracts of testis or spleen from adults. Lower molecular weight bands (around 75 kDa) are independent of tissue origin and are presumably from immunoglobulin and other proteins in the IP antiserum. c) RT-qPCR analysis of whole testis RNA extracted from 6, 10, 12, 16 dpp juvenile animals. Three different primer pairs were used to amplify *Ccnb3*, and three cohorts of littermates were analyzed. The plotted values were the expression levels of *Ccnb3* normalized to *B2M*. The red lines are means across primer pairs. d) IP-WB for cyclin B3 in extracts from testes of juvenile mice at 6 or 16 dpp. The dashed line indicates where the blot image was spliced to remove irrelevant lanes.

To detect endogenous cyclin B3, we used immunoprecipitation and western blotting (IP-WB). Two of the antibodies behaved well for this purpose: antibody #18 (hereafter mAb^N^) raised against a more N-terminal peptide (aa 110-123) and antibody #5 (hereafter mAb^C^) raised against a more C-terminally located peptide (aa 1067-1080) (**Fig 1a**). IP-WB with these antibodies detected endogenous cyclin B3 from wild-type testis extracts but not spleen tissue extracts (**Fig 1b**). The protein migrated as a doublet at ~150 kDa. The slower-migrating band may represent post-translational modification.

To gain further insights into *Ccnb3* expression, we collected testes from wild-type males during the first wave of meiosis at 6, 10, 12, and 16 days postpartum (dpp) and analyzed mRNA levels by reverse-transcription quantitative PCR (RT-qPCR). *Ccnb3* expression was at minimal or background levels at 6 dpp, when seminiferous tubules contain only somatic cells and spermatogonia (the first wave of spermatogenesis begins around 7 dpp in mice (Bellvé et al. 1977)). *Ccnb3* signal was readily detected at 10 dpp and was highest at 12 dpp (**Fig 1c**), when seminiferous tubules are mostly populated with leptotene and zygotene cells (Bellvé et al. 1977). This result agrees with a previous study (Nguyen et al. 2002) and published testis RNA-seq data (Margolin et al. 2014).

Cyclin B3 protein was detected in testis extracts from 16-dpp but not 6-dpp males, confirming that cyclin B3 protein is attributable to spermatocytes (**Fig 1d**). Unfortunately, we have been unable to detect the protein by immunofluorescent staining of meiotic spreads or squashes (data not shown). Nevertheless, these experiments provide the first direct confirmation of the presence of cyclin B3 protein in early meiotic cells.

### Targeted mutation of Ccnb3

To address whether cyclin B3 has a role during spermatogenesis, we generated endonuclease mediated (em) mutations by CRISPR-Cas9-mediated genome editing. We used a guide RNA to target the 3’ end of the 2.7-kb long exon 7. Among the founder animals, several out-of-frame alleles were identified (**Fig 2a**). We focused on the *Ccnb3^em1^* allele (hereafter *Ccnb3*^−^), which has a 14-bp deletion causing a frameshift and premature stop codon upstream of the cyclin box, which is encoded in exons 9–13. We reported elsewhere that the mutant mice are viable with no apparent somatic defects (Karasu et al. 2019).

**Figure 2:**
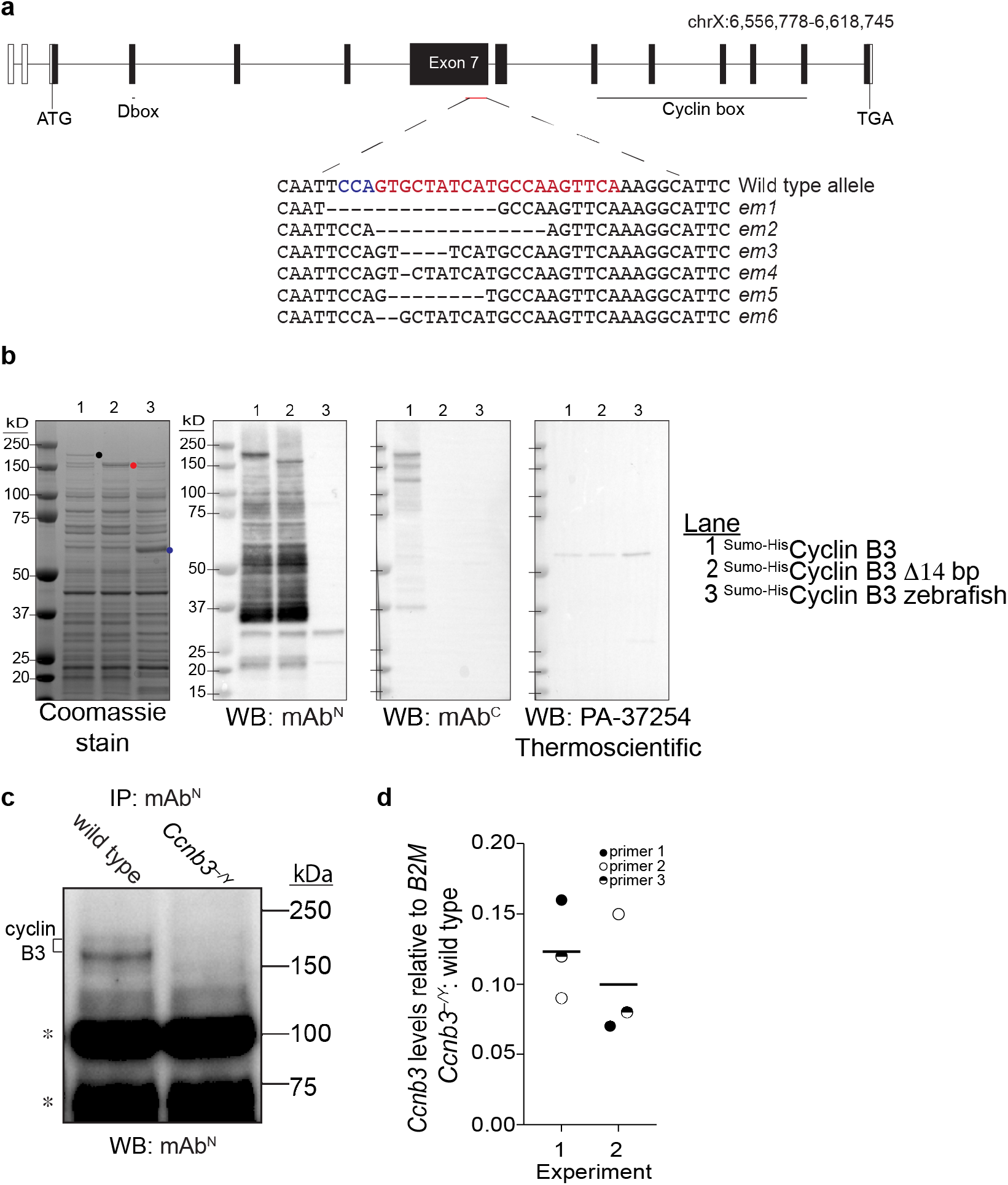
CRISPR/Cas9-mediated Ccnb3 allele generation and confirmation. a) *Ccnb3* gene diagram. Open boxes show 5’ and 3’untranslated regions and filled boxes show the coding regions. The CRISPR-Cas9 targeted position is indicated with protospacer adjacent motif (PAM) in blue and single guide RNA sequence in red. Sequences are shown for six *em* alleles recovered. The *em1* allele was characterized in this study and Karasu et al. 2019. b) Expression and detection of recombinant cyclin B3 proteins. Constructs contained either full-length coding sequence or the 14-bp deletion of the *em1* allele fused to the Sumo-His6 tag (Wasmuth and Lima 2012). Zebrafish cyclin B3 was used as a negative control for detection by WB. Proteins were expressed in *E. coli* and equal aliquots of soluble extracts were fractionated on SDS-PAGE and stained with Coomassie blue or analyzed by WB with the indicated antibody. Dots indicate positions of the largest detected proteins encoded by each construct. Lower molecular weight bands detected by WB with mAb^N^ or mAb^C^ presumably areproteolytic fragments and/or products of premature translation termination. c) IP-WB for cyclin B3. Whole testis extracts from *Ccnb3^−/Y^* and wild-type adults were analyzed using mAb^N^. Note that IP-WB using only this antibody is less sensitive and gives higher background than when combining mAb^N^ with mAb^C^ (**Fig 1b**). Asterisks indicate lower molecular weight bands independent of genotype, possibly from immunoglobulin and other contaminants in the IP elute. d) RT-qPCR analysis of whole testis RNA extracted from *Ccnb3^−/Y^* and wild-type adults. The plotted values represent the fold difference between mutant and wild type for *Ccnb3* levels normalized to *B2M*. Three different pairs of *Ccnb3* primers were used. Two independent experiments were performed, each containing a separate mutant–wild type pair. Lines indicate means for each experiment.

In principle, the *Ccnb3*^−^ allele could produce a C-terminally truncated protein of 1090 aa (~123 kDa). When recombinant cyclin B3 was expressed in *E. coli*, blotting with mAb^N^ detected both full length protein and the shorter form expressed from a construct mimicking the *em1* allele (**Fig 2b**). Full length protein but not the truncated form could be detected with mAb^C^ (**Fig 2b**), as expected because the deletion mutation destroys the epitope recognized by this antibody (**Fig 1a**). Zebrafish cyclin B3 was not detected with either antibody raised against the mouse protein, confirming specificity (**Fig 2b**). Interestingly, a commercial anti-cyclin B3 antibody used in a recent study (Tang et al. 2018) failed to detect the recombinant proteins under these conditions (**Fig 2b**, right panel), suggesting that caution is warranted when using this reagent.

To address whether the C-terminally truncated protein was present in *Ccnb3^−/Y^* spermatocytes, we performed IP-WB experiments using mAb^N^. Testis extracts from *Ccnb3^−/Y^* mutants failed to yield signal above background for either the full length or truncated product (**Fig 2c**). We cannot exclude the possibility that low levels of the truncated protein are present, but these results suggest that the truncated form is unstable *in vivo* if it is made at all. We could not test whether a C-terminal portion of the protein downstream of the mutation is expressed because of lack of a suitable antibody.

Analysis of *Ccnb3* mRNA levels by RT-qPCR showed an ~10-fold decrease in *Ccnb3^−/Y^* mutants compared to wild-type littermates (**Fig 2d**). This decrease may be caused by nonsense-mediated decay and provides further support that the *em1* allele is at least a severe loss-of-function allele.

### Ccnb3^−/Y^ males do not show gross defects in spermatogenesis or meiotic prophase progression

Problems during spermatogenesis typically lead to sterility or reduced fertility, reduced testis weight, and/or seminiferous tubule abnormalities (de Rooij and de Boer 2003). To address whether *Ccnb3^-/Y^* males are fertile, males were placed in cages with wild-type females for two months. *Ccnb3^−/Y^* males and their wild-type littermates sired similar numbers of progeny (totals during this breeding period of 30 and 27 pups, respectively, sired from two females each). Furthermore, adult *Ccnb3^−/Y^* males were indistinguishable from their wild-type littermates for testis/body weight ratio (**Fig 3a**, p = 0.14, Student’s t-test) and in histological appearance of seminiferous tubule sections (**Fig 3b**). Seminiferous tubules from *Ccnb3^−/Y^* contained the full array of spermatogenic cells including spermatogonia, spermatocytes, and round and elongated spermatids (**Fig 3b**). Thus, spermatogenesis is not grossly impaired in the absence of *Ccnb3*.

**Figure 3:**
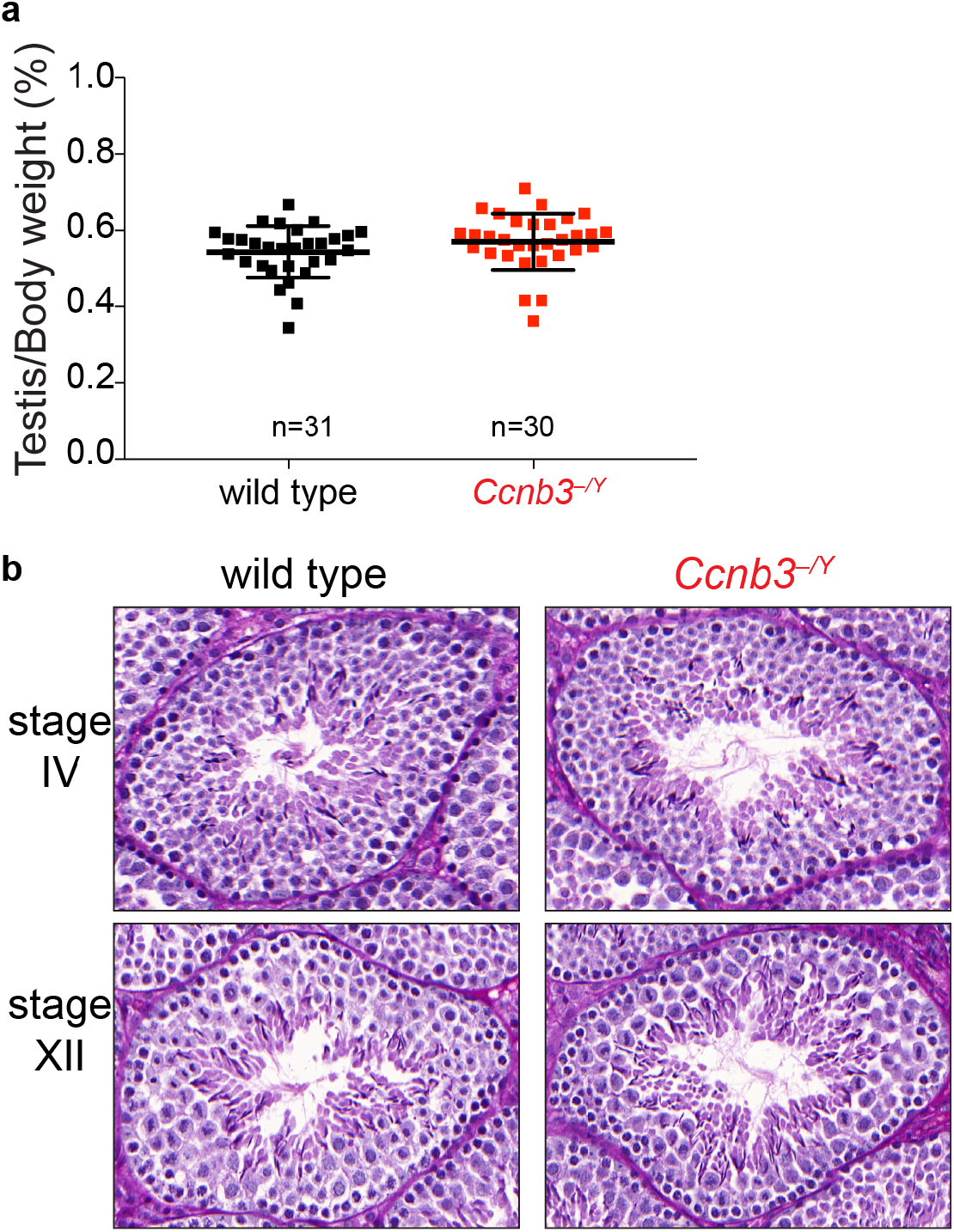
Genetic ablation of Ccnb3 did not affect testis size or seminiferous tubule structure. a) Testis-to-body weight ratios (percent) for adult mice. Error bars indicate means ± standard deviation: 0.54 ± 0.068 and 0.57 ± 0.070 for wild type and *Ccnb3^−/Y^*, respectively. b) Representative periodic acid Schiff stained seminiferous testis sections from adult mice. Stages IV and XII are the stages most commonly associated with apoptosis of spermatocytes encountering meiotic defects (Ahmed and de Rooij 2009).

To test for more subtle meiotic defects, we immunostained meiotic chromosome spreads. In meiotic prophase, cells can be classified into four different cytological stages; leptonema, zygonema, pachynema and diplonema. The transition between each stage can be followed by the changes in the staining pattern of components of the synaptonemal complex (SC) (de Vries et al. 2005; Fraune et al. 2012). SYCP3, a component of the axial elements of the SC, appears as dots or short patches in leptonema. During zygonema, SYCP3-containing axes elongate and SYCP1, which forms the transverse filaments of the SC, starts to appear. The SC is present between homologs along their entire length during pachynema, then the SC disassembles in diplonema. We followed SC formation and disassembly in *Ccnb3^−/Y^* males and their wild type littermates by staining chromosome spreads with antibodies to SYCP3 and SYCP1. We did not observe any aberrant SC formation or defects in disassembly (**Fig 4a**).

**Figure 4:**
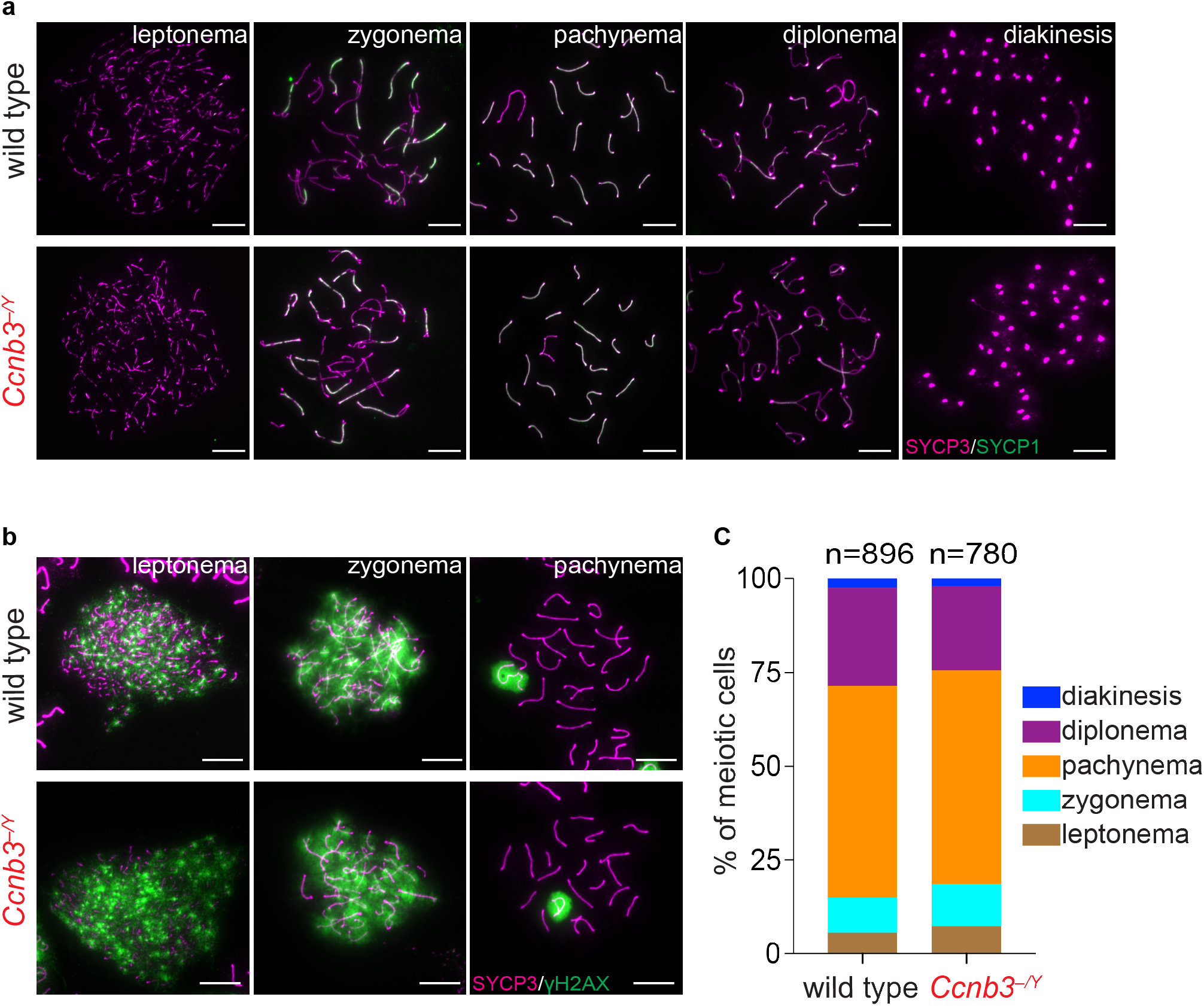
Meiotic Progression is normal in Ccnb3^−/Y^ males. a) Representative images of meiotic prophase stages. Meiotic spreads from wild type and *Ccnb3^−/Y^* males were stained with SYCP3 and SYCP1 antibodies. Scale bars represent 10 μm. b) Representative images of meiotic spreads stained with SYCP3 and γH2AX antibodies. Scale bars represent 10 μm. c) Distribution of meiotic prophase stages. Two mice for each genotype were analyzed. Meiotic cells are defined as follows. Leptonema: short patches of SYCP3 and γH2AX staining on autosomes;zygonema: some synapsed homolog stretches and γH2AX staining on autosomes; pachynema: fully synapsed homologs and γH2AX staining only on sex body; diplonema: desynapsed homologs and γH2AX staining only on sex body; diakinesis: SYCP3 on centromeres and no γH2AX staining.

### Ccnb3^−/Y^ spermatocytes are recombination proficient

We used cytological markers to evaluate the formation and repair of meiotic DSBs. γH2AX is a phosphorylated form of the histone variant H2AX that accumulates upon formation of DSBs (Mahadevaiah et al. 2001). In wild-type mice, γH2AX appears during leptonema then disappears from the autosomes as DSBs are repaired, leaving γH2AX signal only in the sex body, a heterochromatic domain containing the sex chromosomes visible during pachynema and diplonema (Mahadevaiah et al. 2001; Barchi et al. 2005). γH2AX appearance and disappearance were similar in *Ccnb3^−/Y^* males and their wild-type littermates (**Fig 4b**). Defects in meiotic DSB repair often manifest as persistent patches of γH2AX along fully synapsed chromosomes in late pachynema and desynapsing chromosomes in diplonema (Roig et al. 2010; Pacheco et al. 2015; Marcet-Ortega et al. 2017). Absence of such patches in *Ccnb3^−/Y^* males (n = 23 cells examined) suggests that DSB repair is completed normally.

We also counted the number of cells at different stages of meiotic prophase using γH2AX and SYCP3 staining (**Fig 4c**). The distribution of cell stages was indistinguishable between *Ccnb3^−/Y^* males and their wild-type littermates (p = 0.95, chi-square test), further confirming that meiotic progression is affected little if at all by disruption of cyclin B3 expression.

We also counted the number of foci of the strand exchange protein, DMC1; these foci are maximal at early-to-mid zygonema and then decrease as DSBs are repaired via homologous recombination (Moens et al. 2002). *Ccnb3^−/Y^* males displayed overall similar numbers and temporal progression of DMC1 foci as their wild-type littermates, except that they had a significant but quantitatively modest elevation in focus numbers in late zygonema and early-to-mid pachynema (**Fig 5a**). Lastly, we counted MLH1 foci, which mark nascent crossover sites from mid-to-late pachynema (Gray and Cohen 2016). The *Ccnb3^−/Y^* males showed a slight increase in MLH1 foci compared to their wild-type littermates (means of 25 foci in wild type vs. 26 foci in *Ccnb3^−/Y^*)(**Fig 5b**). Overall, these data suggest that *Ccnb3* is largely if not completely dispensable for initiation and completion of meiotic recombination, although it may contribute modestly to control of the amount or timing of recombination (see Discussion).

**Figure 5:**
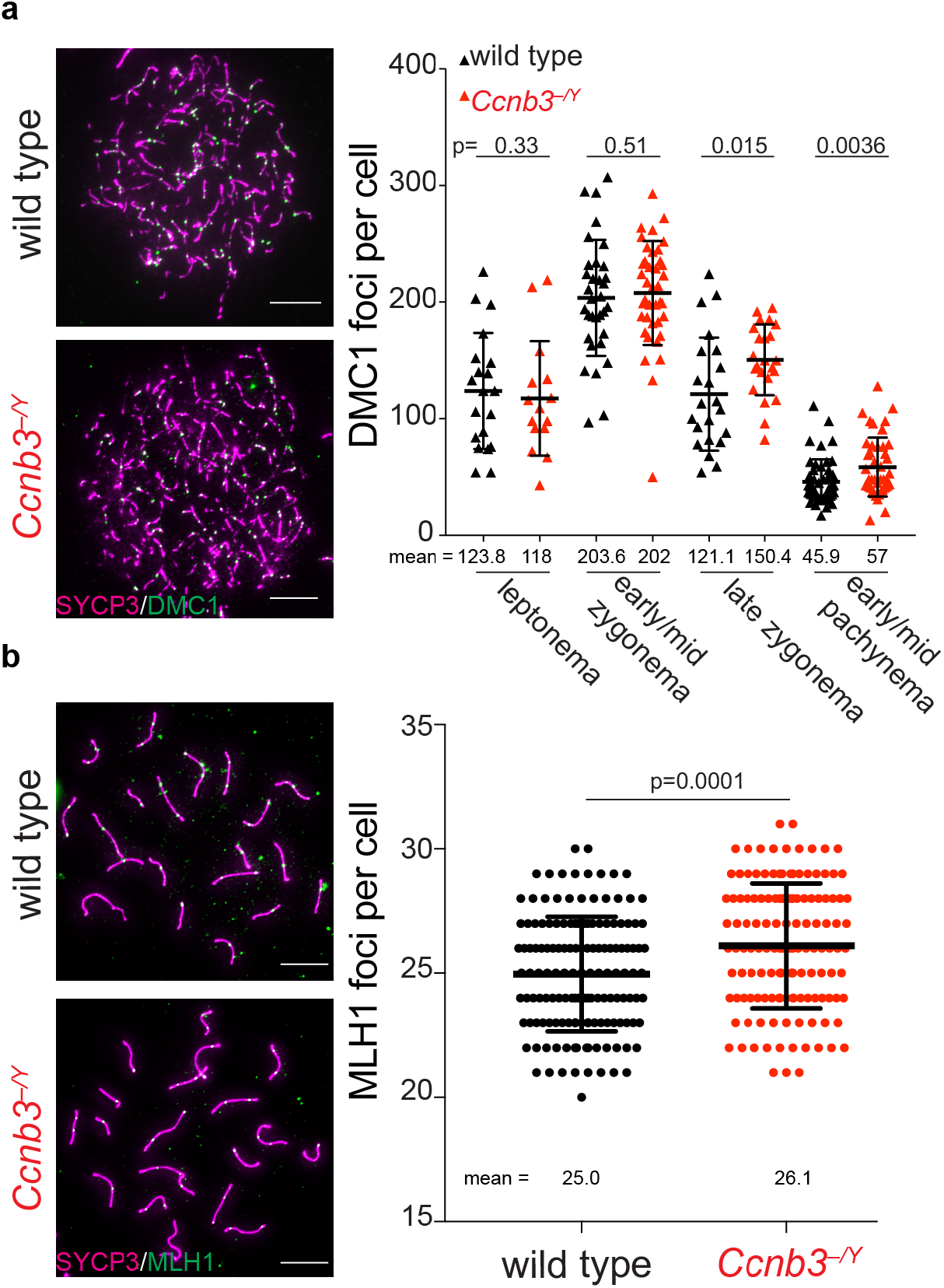
DMC1 and MLH1 focus numbers in Ccnb3^−/Y^ males. a) Representative images (left) and quantification (right) of meiotic spreads stained with DMC1 and SYCP3 antibodies. Scale bars represent 10 μm. b) Representative images (left) and quantifications (right) of meiotic spreads stained with MLH1 and SYCP3 antibodies. Scale bars represent 10 μm. Error bars indicate means ± standard deviation; p values are from Student’s t tests.

### Evaluating potential redundancy with other cyclins

During spermatogenesis, approximately thirty cyclins and cyclin-related genes are present in different cell populations (Malumbres and Barbacid 2005; Malumbres and Barbacid 2009; Margolin et al. 2014), raising the possibility of redundancy between *Ccnb3* and other cyclins.

For example, deletion of *Ccne2* (encoding cyclin E2) causes upregulation of *Ccne1* mRNA and cyclin E1 protein levels, which compensates for the absence of *Ccne2* during spermatogenesis (Martinerie et al. 2014). Likewise, *Ccne2* is upregulated in a *Ccne1* mutant, which compensates for the absence of *Ccne1* (Martinerie et al. 2014). To address whether a similar phenomenon occurs in *Ccnb3^−/Y^* males, we quantified testis mRNA levels of a panel of cyclins by RT-qPCR. We found that *Ccna2* levels were decreased by ~30% whereas others were unchanged (**Fig 6a**). Importantly no upregulation was detected among the tested cyclins. While these results do not rule out the possibility that one of these cyclins exerts a redundant role with *Ccnb3*, the findings suggest that compensatory misregulation such as seen between *Ccne1* and *Ccne2* is not responsible for the lack of meiotic defect in cyclin B3-deficient spermatocytes.

**Figure 6:**
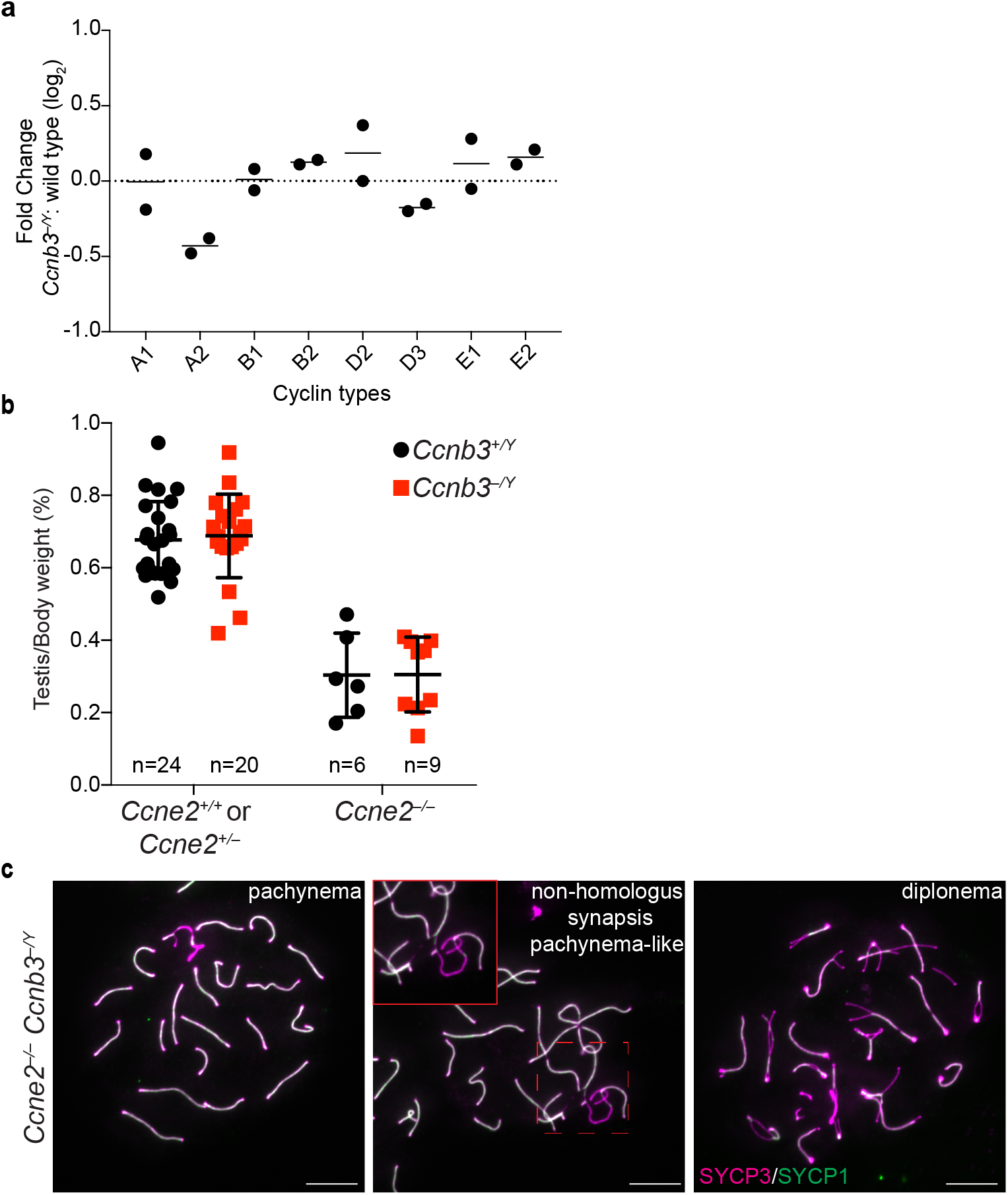
Testing for redundancy with other cyclins. a) RT-qPCR analysis of whole testis RNA extracted from *Ccnb3^−/Y^* and wild-type adults. The plotted values represent the log_2_ fold difference between mutant and wild type for levels of the indicated cyclin mRNA normalized to *B2M*. Two independent experiments were conducted, each containing a separate mutant–wild type pair. Black lines represent the means. b) Testis-to-body weight ratios (percent) for adult mice. Error bars indicate means ± standard deviation for *Ccne2*^+/+^ or *Ccne2^+/−^ Ccnb3^+/Y^*: 0.68 ± 0.11; *Ccne2^+/+^* or *Ccne2^+/−^ Ccnb3^−/Y^*: 0.69 ± 0.14; *Ccne2^−/−^ Ccnb3^+/Y^*: 0.30 ± 0.10; *Ccne2^−/−^ Ccnb3^−/Y^* : 0.30 ± 0.10. c) Representative images of meiotic spreads *Ccne2^−/−^, Ccnb3^−/Y^* spermatocytes stained with SYCP3 and SYCP1 antibodies. Inset shows an example of abnormal synapsis between homologs. Scale bars represent 10 μm.

We next considered the possibility that E-type cyclins might substitute for *Ccnb3* because they are present during overlapping portions of meiotic prophase, particularly *Ccne2* (Martinerie et al. 2014). Testis size is reduced in *Ccne2^−/−^* mice because spermatocytes arrest in diplonema, with a fraction of cells displaying abnormal synapsis during pachynema (Geng et al. 2003; Martinerie et al. 2014). In *Ccne1^−/−^ Ccne2^−/−^* double mutant mice, spermatocytes arrest in a pachytene-like stage with severe pairing and synapsis defects (Martinerie et al. 2014). We asked whether the combined deletion of *Ccne2* and *Ccnb3* would lead to synergistic defects in spermatogenesis. However, introducing the *Ccnb3* mutation did not further exacerbate the reduced testis size in *Ccne2^−/−^* mutants (**Fig 6b**, p ≥0.98, Student’s t-test). Staining meiotic spreads for SYCP1 and SYCP3 (**Fig 6c**) revealed examples of synaptic anomalies in *Ccne2^−/−^ Ccnb3^−/Y^* spermatocytes as previously reported for *Ccne2^−/−^* mutants, and also showed that double mutant spermatocytes could progress through pachynema and reach diplonema, as in *Ccne2^−/−^* (Martinerie et al. 2014). These results indicate that the mutation of *Ccnb3* has no additive effect on the phenotype of *Ccne2* mutants, suggesting in turn that cyclin E2 does not compensate for the absence of cyclin B3.

### Ccnb3^−/Y^ spermatocytes do not show gross abnormality in the metaphase-to-anaphase I transition

Similar to spermatocytes, oocytes deficient for cyclin B3 show no apparent defects in meiotic prophase I (Karasu et al. 2019; Li et al. 2019). However, cyclin B3 is essential for the metaphase-to-anaphase I transition in oocytes. *Ccnb3^−/−^* oocytes arrest in metaphase I with compromised activity of the anaphase promoting complex/cyclosome (APC/C), which leads to inefficient degradation of the APC/C substrates cyclin B1 and securin and no active separase(Karasu et al. 2019; Li et al. 2019). Our results thus far showed no evidence of metaphase arrest in cyclin B3-deficient spermatocytes, as judged by the normal size of *Ccnb3^−/Y^* testes (**Fig 3a**) and normal appearance of stage XII tubules (**Fig 3b**; tubules of this type contain meiotically dividing spermatocytes). However, to more rigorously assess whether *Ccnb3^−/Y^* spermatocytes experience a delayed metaphase-to-anaphase transition, we determined the number of stage XII tubules. Defects in the MI-to-MII transition can elevate the fraction of stage XII tubules (Papanikos et al. 2018). Testis sections were stained with antibodies against histone H3 phoshorylated on serine 10 (pH3) (Jain et al. 2018) (**Fig 7a**). The percentage of tubules that were at stage XII was similar between wild type and *Ccnb3^−/Y^*, suggesting that mutant spermatocytes do not reside longer in metaphase I during spermatogenesis (**Fig 7b**). These results lead us to infer that APC/C activity is likely not substantially compromised in spermatocytes lacking cyclin B3, unlike in oocytes.

**Figure 7:**
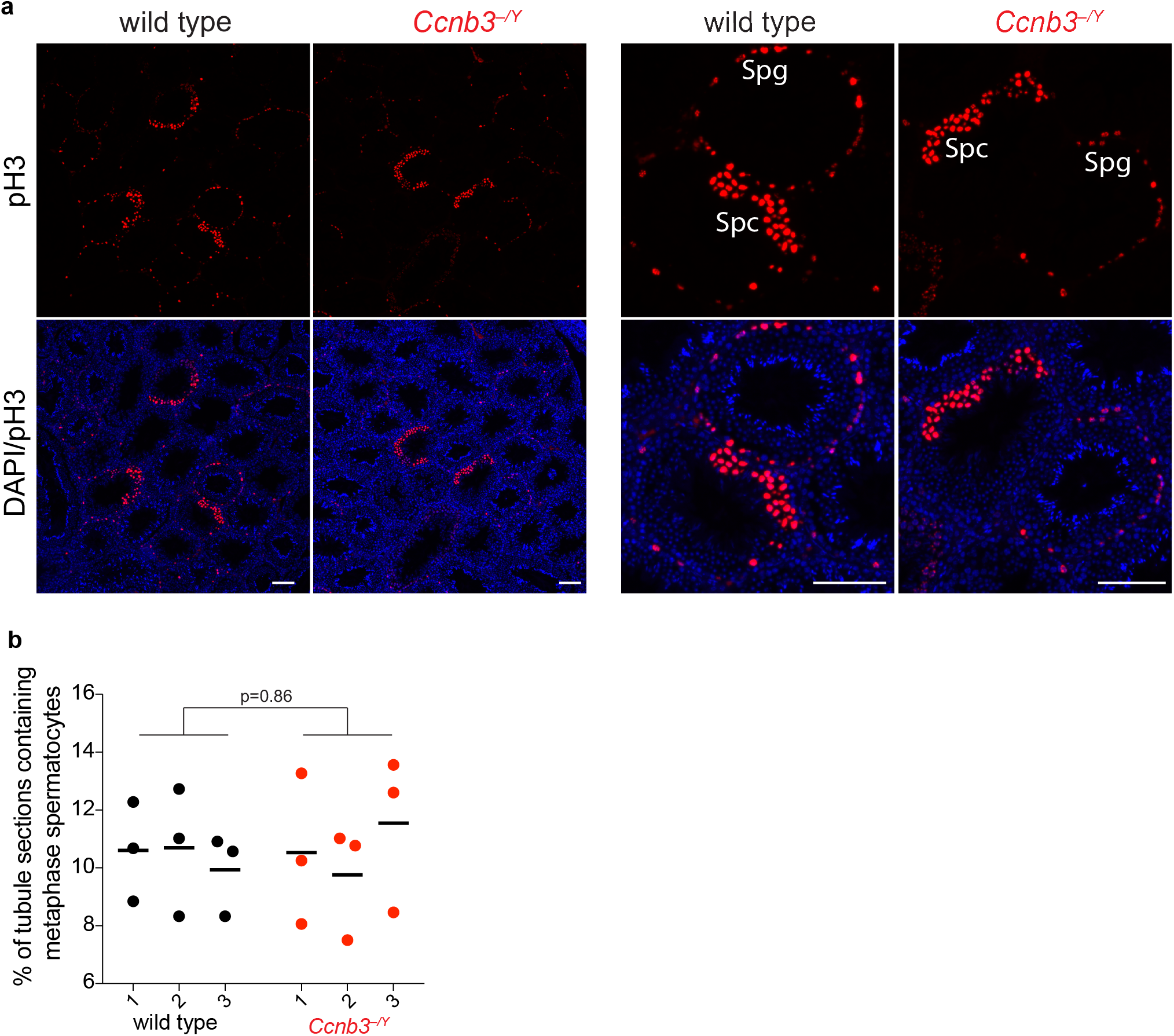
No apparent accumulation of metaphase spermatocytes in Ccnb3^−/Y^ males. a) Representative pH3-stained testis sections from wild type and *Ccnb3^−/Y^* (lower magnification views at left; higher magnification at right). Anti-pH3 also stains mitotically dividing spermatogonia, which are easily recognized because they are located at the periphery of the tubules; these were excluded from the counting. Spg: spermatogonia, Spc: spermatocytes. Scale bar represents 100 μm. b) Quantification of tubules containing meiotically dividing cells. Each point is the value from one testis section. Three sections were scored from each of three different animals per genotype. Lines indicate means for each animal. The p value is from a Student’s t test performed on data pooled for each genotype.

## Discussion

Despite prominent, regulated expression during spermatogenesis, *Ccnb3*-deficient males are fully fertile and exhibit no gross meiotic abnormalities. While we cannot exclude the possibility that the *em1* allele retains partial *Ccnb3* function, our molecular characterization of mRNA and protein expression in testes (this study) and the strong metaphase I arrest in homozygous mutant oocytes (Karasu et al. 2019) show that this allele is at least a substantial loss-of-function mutation. Furthermore, an independently generated *Ccnb3* mutation that similarly causes oocyte arrest was also reported to support normal male fertility (Li et al. 2019). Taken together, these findings strongly suggest that cyclin B3 is dispensable for male meiosis. The basis of the extreme sexual dimorphism in the requirement for cyclin B3 is thus far unclear. Null *Ccnb3* mutations in *Drosophila melanogaster* also do not affect male fertility (Jacobs et al. 1998), although differences in meiosis in male flies such as lack of recombination (McKee et al. 2012) raised the possibility that the genetic requirement could have been different in mice.

Based on its expression timing, we originally speculated that *Ccnb3^−/Y^* males might have defects in processes important for meiotic recombination, such as DSB formation. If so, we would have expected a similar phenotype to mice that lack or have reduced activity of essential DSB proteins, including the *Spo11, Mei4, Mei1, Iho1, Top6bl* knock out animals (Baudat et al. 2000; Libby et al. 2002; Kumar et al. 2010; Kauppi et al. 2011; Kauppi et al. 2013; Robert et al. 2016; Stanzione et al. 2016). Another possibility might have been that *Ccnb3* contributes to later events, in which case we might have expected phenotypes similar to animals lacking *Ccne1* or *Cdk2* (Geng et al. 2003; Martinerie et al. 2014). However, *Ccnb3^−/Y^* mice did not show any obvious defects associated with spermatogenesis, although we did observe a modest elevation of foci of RAD51 and MLH1. It is possible that this elevation is not biologically meaningful. For example, there could be slight differences in strain background between *Ccnb3^−/Y^* mice and the controls because of incomplete backcrossing, or there could be small differences between the samples in terms of enrichment for slightly later or slightly earlier cells within the scored substages of meiotic prophase. However, it is also possible that cyclin B3 contributes modestly to regulating how quickly recombination is completed. Alternatively, absence of cyclin B3 may allow cells to continue forming DSBs past the time they normally would have stopped. Importantly, however, the absence of γH2AX patches in late pachynema or diplonema and the apparently normal meiotic progression of *Ccnb3^−/Y^* spermatocytes strongly indicate that cyclin B3 is largely if not completely dispensable for completion of recombination.

It is intriguing that a gene specifically expressed in testis and whose temporal expression in male mice is confined to meiotic prophase appears not to contribute to meiosis or spermatogenesis in any substantive manner. However, a recent study reported the generation of knock-out mutations for 54 testis-specific genes, all of which yielded fertile mice (Miyata et al. 2016). It could be that all or most of these genes are not important at all for reproductive success, but an interesting alternative is that their functions are context dependent. It should be noted that nearly all experiments to date have been conducted with inbred laboratory mice that are not under stress or reproductive competition. These experiments may therefore fail to capture biological functions that would manifest in more natural conditions. For example, one of the 54 genes knocked out by Miyata and colleagues was the polycystin family receptor for egg jelly, *Pkdrej*. *Pkdrej*-deficient males were as fertile as their wild-type littermates in unrestricted mating trials, but they performed poorly when competing with wild type in sequential mating trials and in artificial insemination of mixed-sperm populations, indicating that *Pkdrej* is important in postcopulatory reproductive selection (Sutton et al. 2008). In light of findings such as this, it may be interesting to query the effects of *Ccnb3* mutations in different strain backgrounds or in more complex mating scenarios that are more akin to reproductive challenges faced in natural environments.

## Materials and Methods

### Animals

Mice in the Keeney lab were maintained and sacrificed for the experiments under regulatory standards approved by the Memorial Sloan Kettering Cancer Center (MSKCC) Institutional Animal Care and Use Committee. Animals were fed regular rodent chow with ad libitum access to food and water.

### Antibody production

Mouse monoclonal antibodies directed against mouse cyclin B3 were produced by Abmart Inc. (Shanghai) using synthetic peptides of the sequences shown in **Fig 1**. Ascites from different immunization clones were first tested by western blotting against recombinant cyclin B3 protein expressed in *E. coli* from the pGKBT7 vector (Clontech). Clones that recognized the recombinant protein were tested for their efficiency in immunoprecipitating recombinant cyclin B3. Hybridoma cell lines were generated for mAb^C^ and mAb^N^ antibodies and were maintained in the Antibody and Bioresource Core Facility at MSKCC. Cell culture supernatants from the cell lines were subjected to protein G purification to enrich monoclonal antibodies and antibodies were stored in phosphate-buffered saline (PBS) at −20°C.

### Immunoprecipitation and western blotting from testis extracts

For mouse testis extracts, dissected testes were placed in an Eppendorf tube, frozen on dry ice, and stored at −80°C. The frozen tissue was suspended in RIPA buffer (50 mM Tris-HCl pH 7.5, 150 mM NaCl, 0.1% SDS, 0.5% sodium deoxycholate, 1% NP40) supplemented with protease inhibitors (Roche Mini tablets). The tissue was disrupted with a plastic pestle and incubated by end-over-end rotation for 15 minutes at 4°C. After brief sonication, samples were centrifuged at 15000 rpm for 15 minutes. The clear lysate was transferred to a new tube and used for immunoprecipitation.

Testis extract was pre-cleared with protein G Dynabeads (Thermofisher) by end-over-end rotation for 1 hr at 4°C. Antibodies were added to pre-cleared lysates and incubated overnight with end-over-end rotation at 4°C. Protein G Dynabeads were added to the tubes and incubate for 1 hr with end-over-end rotation at 4°C. Beads were washed three times with RIPA buffer, resuspended in 1x NuPAGE LDS sample buffer (Invitrogen) with 50 mM DTT, and boiled 5 minutes to elute immunoprecipitated proteins.

Immunoprecipitated samples were separated on 4–12% Bis-Tris NuPAGE precast gels (Life Technologies) at 150 V for 70 minutes. Proteins were transferred to polyvinylidene difluoride (PVDF) membranes by wet transfer method in Tris-glycine-20% methanol at 120 V for 40 minutes at 4°C. Membranes were blocked with 5% non-fat milk in PBS-0.1% Tween (PBS-T) for 30 minutes at room temperature on an orbital shaker. Blocked membranes were incubated with primary antibodies 1 hr at room temperature or overnight at 4°C. Membranes were washed with PBS-T for 30 minutes at room temperature, then incubated with horseradish peroxidase (HRP)-conjugated secondary antibodies for 1 hr at room temperature. Membranes were washed with PBS-T for 15 minutes and the signal was developed by ECL Plus (Perkin Elmer) or ECL Prime (GE Healthcare).

### CRISPR-Cas9 mediated gene targeting and mouse genotyping

*Ccnb3* mutations were generated by the MSKCC Mouse Genetics Core Facility. Exon 7 was targeted by guide RNA (C40-TGAACTTGGCATGATAGCAC). Guide RNA was cloned into pDR274 vector for *in vitro* transcription. *In vitro* transcribed guide RNA (10 ng/μl) and Cas9 (20 ng/μl) were microinjected using conventional techniques (Romanienko et al. 2016) into pronuclei of CBA/J × C57BL/6J F2 hybrid zygotes generated by crossing CBAB6F1/J hybrid females with C57BL/6J males.

Genomic DNA from founder animals was subjected to PCR by using primers CCN3B-A and CCN3B-B (all primer sequences are in **Supplemental Table S1**) under the following conditions: 3 minutes at 94°C; then 35 cycles of 15 seconds at 94°C, 90 seconds at 68°C; then a final extension for 3 minutes at 68°C. T7 endonuclease I digestion was performed to identify the animals carrying insertion/deletion mutations (indels) (Guschin et al. 2010). Because males have only one copy of the X-linked *Ccnb3* gene, the PCR amplification and T7 assay on male founders was performed in the presence of wild-type genomic DNA.

To define the molecular nature of indels, genomic DNA from T7-positive animals was amplified using the primers indicated above. Amplification products were used for TA Cloning (TA Clonining™ Kit with pCR™ 2.1 vector, Invitrogen). Ten white bacterial colonies were selected and inserts were sequenced. The *em1* allele, an out of frame deletion, was chosen to generate the *Ccnb3^em1sky^* line. After two rounds of backcrossing to C57BL/6J, animals were interbred to generate homozygous and heterozygous female animals and hemizygous male animals.

Primers for genotyping were GT4-F and GT4-R. The PCR was performed under following conditions: 2 minutes at 94°C; then 35 cycles of 20 seconds at 94°C, 30 seconds at 54°C and 30 seconds at 72°C; then a final extension 3 minutes at 72°C. The amplification product (202 bp) was subjected to BsrI enzyme digestion (NEB). The wild-type sequence is cut to yield two fragments of 103 and 99 bp; the BsrI site is lost in the *em1* allele.

For cyclin E genotyping, the published protocol was followed (Martinerie et al. 2014). Briefly, the following primers were mixed and used for PCR: E2-L, E2-G, and EN-3. The PCR was performed as follows: 2 minutes at 94°C; then 35 cycles of 20 seconds at 94°C, 30 seconds at 54°C and 30 seconds at 72°C; then a final extension 3 minutes at 72°C. The PCR products are 400 bp for the wild-type allele and 300 bp for the mutant allele. Further genotyping was performed by PCR with primers E2-3 and E2-4 using the following conditions: 2 minutes at 94°C; then 35 cycles of 20 seconds at 94°C, 30 seconds at 54°C and 30 seconds at 72°C; then a final extension 3 minutes at 72°C. A 223 bp PCR product is recovered from wild type; absence of PCR product indicates the mutant.

### Spermatocyte meiotic spread preparation

Primary and secondary antibodies are listed in **Supplemental Table S2**. Testes were dissected and deposited after removal of the tunica albuginea in 50 ml Falcon tubes containing 2 ml TIM buffer (104 mM NaCl, 45 mM KCl, 1.2 mM MgSO_4_, 0.6 mM KH_2_PO_4_, 6.0 mM sodium lactate, 1.0 mM sodium pyruvate, 0.1% glucose). 200 μl collagenase (20 mg/ml in TIM buffer) was added, and left shaking at 550 rpm for 55 minutes at 32°C. After incubation, TIM buffer was added to a final volume of 15 ml, followed by centrifugation for 1 min at 600 rpm at room temperature. Separated tubules were washed 3 times in TIM buffer. Then, separated tubules were resuspended in 2 ml TIM, with 200 μl trypsin (7 mg/ml in TIM) and 20 μl DNase I (400 μg/ml in TIM buffer) and incubated for 15 minutes at 32°C at 550 rpm in a thermomixer. 500 μl trypsin inhibitor (20 mg/ml in TIM) and 50 μl DNase I solution were added and mixed. A wide mouthed plastic Pasteur pipette or P1000 pipette was used to disperse the tissue further by pipetting up and down for 2 minutes. Cells were passed through a 70-μm cell strainer into a new 50 ml Falcon tube. TIM was added to a final volume of 15 ml and mixed. Cells were centrifuged for 5 minutes at 1200 rpm. Supernatant was removed, 15 μl DNase I solution was added and gently mixed, followed by 15 ml TIM. Washing with TIM and resuspension in the presence of DNase I was repeated 3 times. Single-cell suspension was pelleted and resuspended in TIM according to original weight (~200 mg in 400 μl). 10 μl of cell suspension was added to 90 μl of 100 mM sucrose solution, flicked three times and incubated for 8 minutes at room temperature.

Superfrost glass slides were divided in two squares by use of Immedge pen. Each square received 100 μl 1% paraformaldehyde (PFA) (freshly dissolved in the presence of NaOH at 65°C, 0.15% Triton, pH 9.3, cleared through 0.22 μm filter) and 45 μl of cell suspension was added per square, swirled three times, and dried in a closed slide box for 3 hours, followed by drying with half-open lid 1.5 hours at room temperature. Slides were washed in a Coplin jar 2 × 3 minutes in milli-Q water on a shaker, 1 × 5 minutes with 0.4%PhotoFlo, air dried and stored in aluminum foil at −80°C.

For staining, slides were washed with PBS and blocked with blocking buffer (PBS with 0.3% bovine serum albumin (BSA)) for 30 minutes at room temperature. Slides were incubated with primary antibodies overnight at 4°C. Slides were washed with PBS-T three times for 10 minutes and then incubated with the secondary antibodies for 1 hr at room temperature. The slides were further washed with PBS-T three times in the dark before air drying and mounting with Vectashield containing 4’,6-diamidino-2-phenylindole (DAPI).

Images of spread spermatocytes were acquired on a Zeiss Axio Observer Z1 Marianas Workstation, equipped with an ORCA-Flash 4.0 camera, illuminated by an X-Cite 120 PC-Q lightsource, with either 63× 1.4 NA oil immersion objective or 100× 1.4 NA oil immersion objective. Marianas Slidebook (Intelligent Imaging Innovations, Denver Colorado) software was used for acquisition.

### Total mRNA extraction, cDNA library generation and RT-qPCR

Testes were dissected and frozen on dry ice. Total mRNA was extracted using RNeasy Plus Mini Kit (QIAGEN, 74134) following the manufacturer’s instructions. Superscript™ III First-Strand Synthesis SuperMix (Invitrogen, 18080400) was used with oligo dT primers to generate testis cDNA, which was diluted 1:10 to be used in RT-qPCR carried out using LightCycler 480 SYBR Green I Master (Roche, 4707516001) under the following conditions: 10 minutes at 95°C; then 45 cycles of 10 seconds at 95°C, 20 seconds at 55°C and 10 seconds at 72°C. Amplification products were detected on the LightCycler 480 II Real-Time PCR instrument (Roche). LightCycler 480 Software was used to quantify products by absolute quantification analysis using the second derivative maximum method. All reactions were done in triplicate and the mean of crossing point (Cp) value were used for the analysis. Cp values were normalized to the value obtained for *B2M* reactions (ΔCp). Then, the differences between knockout and wild-type samples were calculated for each primer set (ΔΔCp) and the fold change (knockout vs wild type) was calculated as 2^−_ΔΔ_Cp^. Primers used for RT-qPCR are listed in **Supplemental Table S1**.

### Plasmids

Mouse *Ccnb3* mRNA was amplified by PCR from whole testis cDNA and cloned into modified pGBKT7 vector to express in yeast. Vectors for expression of cyclin B3 (wild type, Δ14 bp mutant or zebrafish) N-terminally tagged with SUMO protein and 6× histidine were generated by cloning into modified pRSF-DUET vector (gift from E. Wasmuth, MSKCC). To generate the Δ14bp mutation, PCR primers that delete the same base pairs that are deleted in the *Ccnb3*^−^ allele were used to amplify pRSF-Duet-*Ccnb3*, and the resulting PCR product was phosphorylated and ligated. Primers are listed in **Supplemental Table S1**.

### Expression of recombinant cyclin B3 in E.coli

The expression vectors described above were used to produce recombinant cyclin B3. Briefly, *E. coli* strain BL21 was transformed with the expression vector and a single colony was picked to inoculate liquid LB overnight at 37°C. The overnight culture was diluted to ~0.04 (OD_600_) and once the OD_600_ had reached ~0.6, the protein expression was induced by addition of 1 mM isopropyl β-D-1-thiogalactopyranoside (IPTG) for 2 hr at 37°C. Cells were harvested by centrifugation at 6000 g, 15 min. All further steps were executed at 4°C. Pellets were resuspended in lysis buffer (50 mM HEPES-NaOH pH 8.0, 300 mM NaCl, 0.1 mM dithiothreitol (DTT), 10% glycerol, and 20 mM imidazole). The sample was sonicated 3 times 10 seconds on-off, 20% duty cycle. The solution was cleared by centrifugation at 15,000 rpm, 20 minutes. The clear protein lysate was mixed with sample loading buffer and analyzed by SDS-PAGE or used for WB.

### Histology

Fixation, tissue processing, and staining were performed as described (Jain et al. 2018). Testes from adult mice were fixed overnight in 4% PFA at 4°C. PFA-fixed tissues were washed for 1 hr in water at 4°C. Fixed tissues were stored in 70% ethanol for up to 5 days prior to embedding in paraffin and sectioning (5 μm for testes). The tissue sections were deparaffinized with EZPrep buffer (Ventana Medical Systems) and antigen retrieval was performed with CC1 buffer (Ventana Medical Systems). Sections were blocked for 30 min with Background Buster solution (Innovex), followed by avidin-biotin blocking for 8 min (Ventana Medical Systems). Periodic acid Schiff (PAS) staining was performed by the MSKCC Molecular Cytology Core Facility using the Autostainer XL (Leica Microsystems, Wetzlar, Germany) automated stainer for PAS with hematoxylin counterstain and coverslips were mounted with Permount (Fisher Scientific). For immunofluorescent staining, slides were deparaffinized with EZPrep buffer (Ventana Medical Systems), antigen retrieval was performed with CC1 buffer (Ventana Medical Systems), and slides were blocked for 30 minutes with background buster solution (Innovex, Richmond, California). For staining phospho-Histone H3 (Ser10), avidin-biotin blocking (Ventana Medical Systems) was performed for 8 minutes. Slides were incubated with the primary antibody (rabbit anti-pH3) for 5 hours, followed by 60 minutes incubation with biotinylated goat anti rabbit antibodies. Streptavidin-HRP D (Ventana Medical Systems) was used for detection followed by incubation with Tyramide Alexa Fluor 594 (Invitrogen, Carlsbad, California). After staining, slides were counterstained with 5 μg/ml, DAPI (Sigma, Damstadt, Germany) for 10 minutes and mounted with coverslips with Mowiol. PAS-stained slides were digitized using Panoramic Flash Slide Scanner (3DHistech, Budapest, Hungary) with a 20× 0.8 NA objective (Carl Zeiss, Jena, Germany). Images were produced and analyzed using the Panoramic Viewer software (3DHistech). Immunofluorescence images for pH3 staining were captured by a Zeiss Axio Observer Z1 Marianas Workstation, equipped with an ORCA-Flash 4.0 camera, illuminated by an X-Cite 120 PC-Q lightsource, either with 10 or 40× 1.4 NA objective. Marianas Slidebook (Intelligent Imaging Innovations, Denver Colorado) software was used for acquisition>

## End Matter

### Author Contributions and Notes

M.E. Karasu performed all of the experiments. S. Keeney provided overall supervision and funding acquisition. Figures were prepared by M.E. Karasu and the manuscript was written by M.E. Karasu and S. Keeney. The authors declare no conflict of interest.

## Acknowledgments

We thank P. Romanienko, W. Mark, J. Ingenito, and J. Giacalone (MSKCC Mouse Genetics Core Facility) for generating mutant mice; F. Weis-Garcia (MSKCC Antibody and Bioresource Core Facility) for monoclonal antibody expression; D. Wolgemuth (Columbia University) for generously sharing *Ccne2^+/−^* animals; C. Claeys Bouuaert (Keeney lab) for critical reading of the manuscript; M. Boekhout and D. Ontoso (Keeney lab) for help with animal handling; N. Fang, M. Turkekul, A. Barlas, and K. Manova-Todorova (MSKCC Molecular Cytology Core Facility) for help with testis histology; and members of Keeney lab for discussion. MSKCC core facilities are supported by NIH cancer center core grant P30 CA008748. This work was supported by the Howard Hughes Medical Institute.

---

**Supplemental Table S1.**
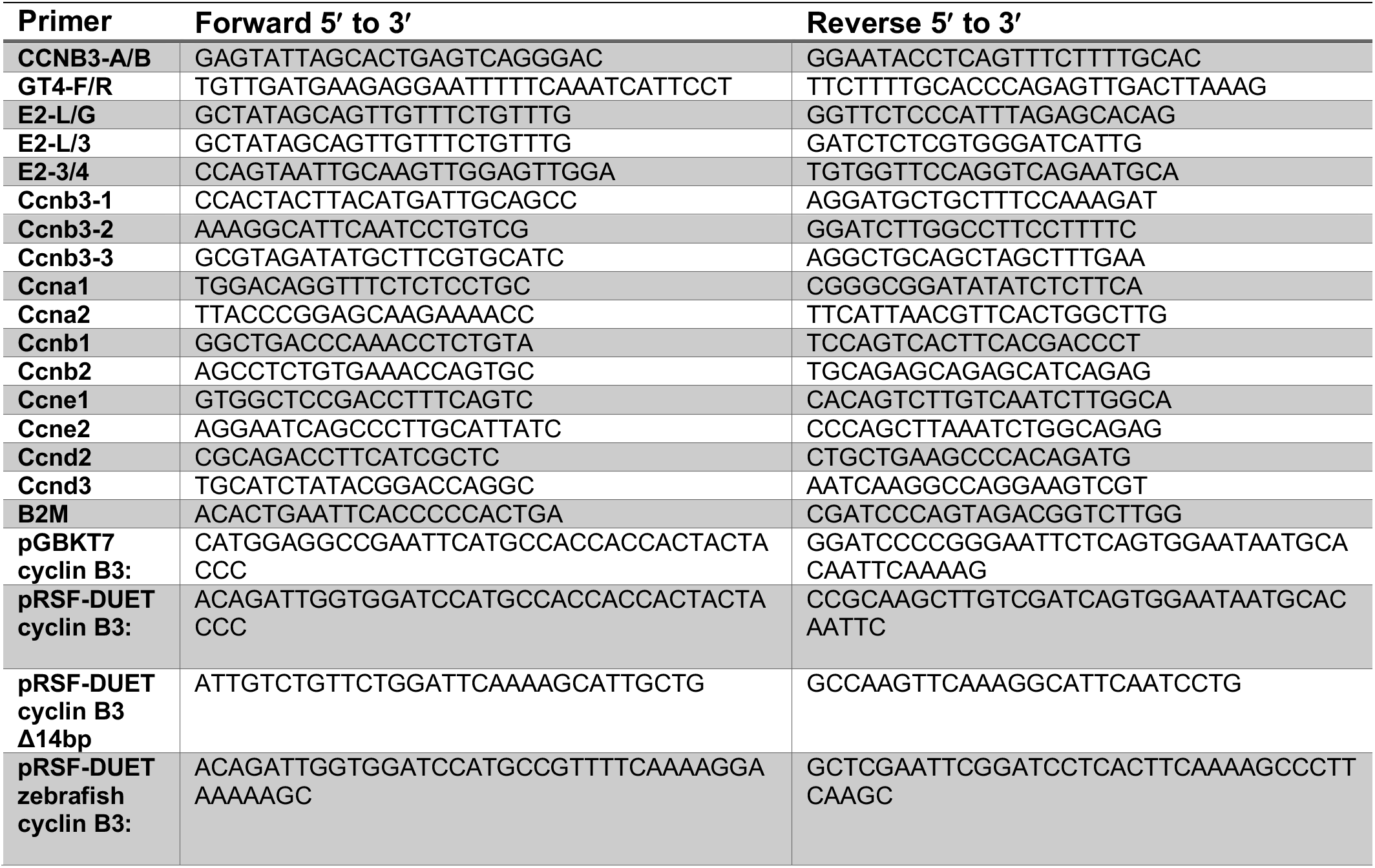
Oligonucleotides

**Supplemental Table S2.** Antibodies

A. Primary Antibodies:
| Target | Amount for IP | Dilution for WB | Dilution for IF/IHC* | Host | Supplier | Catalog |
| --- | --- | --- | --- | --- | --- | --- |
| Cyclin B3-mAB<sup>N</sup> | 10 µg | 1:1000-3000 | N/A | Mouse | Abmart | C517 |
| Cyclin B3-mAB<sup>C</sup> | 2 µg | 1:1000-3000 | N/A | Mouse | Abmart | C500 |
| Cyclin B3 | N/A | 1:1000 | N/A | Rabbit | Thermoscientific | PA5-37254 |
| SYCP3 | N/A | N/A | 1:200 | Mouse | Santa Cruz | sc-74569 |
| γH2AX | N/A | N/A | 1:500 | Rabbit | Santa Cruz | sc-101696 |
| DMC1 | N/A | N/A | 1:100 | Rabbit | Santa Cruz | sc-22768 |
| SYCP1 | N/A | N/A | 1:100 | Rabbit | Abcam | ab15090 |
| MLH1 | N/A | N/A | 1:25 | Rabbit | Calbiochem | pc56 |
| pH3 | N/A | N/A | 1 µg/ml | Rabbit | Upstate | 06-570 |

B. Secondary Antibodies:
| Target | Dilution for WB | Dilution for IF/IHC* | Host | Supplier | Catalog |
| --- | --- | --- | --- | --- | --- |
| anti-mouse | N/A | 1:400 | Donkey | Life Technologies | A21203 |
| anti-rabbit | N/A | 1:400 | Donkey | Thermofisher Scientific | R37118 |
| anti-mouse | 1:10000 |  | Goat | Biorad | 1721011 |
| anti-rabbit | 1:10000 |  | Goat | Biorad | 1706515 |
| anti-rabbit | N/A | 1:200 | Goat | Vector Labs | BA-1000 |

\* IF/IHC, immunofluorescence or immunohistochemistry.

